# Novel *Kras*-mutant murine models of non-small cell lung cancer possessing co-occurring oncogenic mutations and increased tumor mutational burden

**DOI:** 10.1101/2020.02.15.950220

**Authors:** Ramin Salehi-Rad, Rui Li, Linh M. Tran, Raymond J. Lim, Jensen Abascal, Milica Momcilovic, Stacy J. Park, Stephanie L. Ong, Zi Ling Huang, Manash Paul, David B. Shackelford, Kostyantyn Krysan, Bin Liu, Steven M. Dubinett

**Affiliations:** Department of Medicine, Division of Pulmonary and Critical Care, David Geffen School of Medicine at UCLA, 10833 Le Conte Avenue, 43-229 CHS, Los Angeles, CA 90025-1690, USA; Department of Medicine, VA Greater Los Angeles Healthcare System, 11301 Wilshire Boulevard, Los Angeles, CA 90073, USA; Department of Molecular and Medical Pharmacology, David Geffen School of Medicine at UCLA, 650 Charles E. Young Drive South, 23-120 CHS, Box 951735, Los Angeles, CA 90095-1735, USA; Department of Pathology and Laboratory Medicine, David Geffen School of Medicine at UCLA, 757 Westwood Plaza, Los Angeles, CA 90095, USA; Jonsson Comprehensive Cancer Center, UCLA, 8-684 Factor Building, Box 951781, Los Angeles, CA 90095-1781, USA

**Keywords:** Mouse cancer models, NSCLC, TMB, KRAS, LKB1, immunotherapy

## Abstract

Despite recent advances in lung cancer immunotherapy, a major obstacle to the progress in the field is the lack of preclinical models that recapitulate the genetic and immunologic complexity of human disease. Conditional genetically engineered mouse models (GEMMs) of non-small cell lung cancer (NSCLC) harbor the common oncogenic mutations of the disease, but these models possess low tumor mutational burden (TMB), which limits their utility in immunotherapy studies. Here, we establish novel *Kras*-mutant murine models of NSCLC bearing common genetic alterations associated with the disease and increased TMB, by *in vitro* exposure of cell lines derived from GEMMs of NSCLC [*Kras*^*G12D*^ (K), *Kras*^*G12D*^*Tp53*^*−/−*^ (KP), *Kras*^*G12D*^*Tp53*^*+/−*^*Lkb1*^*−/−*^ (KPL)] to the alkylating agent *N*-methyl-*N*-nitrosourea (MNU). Increased TMB was associated with enhanced anti-tumor T cell responses and improved anti-PD-1 efficacy in syngeneic models, across all genetic backgrounds. However, anti-PD-1 efficacy was comparatively modest in the KPL cell lines with increased TMB, which possessed a distinct immunosuppressed tumor microenvironment (TME) primarily composed of granulocytic myeloid-derived suppressor cells (G-MDSCs). This phenotype is consistent with findings in human NSCLC where LKB1 loss is a driver of primary resistance to PD-1 blockade. In summary, these novel *Kras*-mutant murine NSCLC models bearing common co-occurring mutations with increased TMB possess clinically relevant TMEs and recapitulate the genetic complexity and therapeutic vulnerabilities of human NSCLC. We anticipate that these immunogenic models will facilitate the development of novel immunotherapies in NSCLC.

## Introduction

Immune checkpoint inhibitors (ICIs) targeting the PD-1/PD-L1 axis have resulted in durable clinical responses and improved survival in NSCLC^1–4^. However, most patients either do not respond to treatment or develop resistance to therapy after an initial response. Favorable responses to ICIs are associated with high TMB, preexisting CD8^+^T cell tumor infiltration, and high baseline PD-L1 expression within the TME^5–7^. An increased number of candidate MHC class-I tumor-neoantigens and a clonal neoantigen burden have also been associated with improved responses to ICIs in NSCLC^8,9^. Furthermore, a recent study demonstrates that anti-tumor responses to PD-1 blockade derive from a distinct repertoire of T cell clones^10^. These results support the hypothesis that the non-synonymous somatic-mutations associated with increased TMB generate tumor-neoantigens which can be recognized by host T cells as non-self^11^. Treatment with ICIs can stimulate neoantigen-specific T cells to mediate tumor regression.

While ICIs have transformed the treatment landscape of NSCLC, a key impediment to progress in the field of lung cancer immunotherapy is the lack of preclinical models that recapitulate the genetic complexity of human malignancy. Human NSCLCs which are frequently associated with tobacco smoking have among the highest mutational burden of all malignancies and commonly possess genomic alterations in oncogenic pathways^12^. *KRAS* mutations are the most prevalent oncogenic drivers in NSCLC and frequently co-occur with mutations in *TP53* and *LKB1*, which define subgroups of patients with distinct biology^13^. Although conditional GEMMs of *Kras*-mutant NSCLC have served as valuable models in elucidating mechanisms of lung tumorigenesis, studies reveal that these models harbor low TMB with few protein-altering mutations^14–16^. As a result, these GEMMs of NSCLC have limited value in the evaluation of host anti-tumor immune responses in preclinical immunotherapy trials^17^. Herein we report novel *Kras*-mutant murine models of NSCLC bearing common oncogenic mutations of the disease and increased TMB that recapitulate the genetic complexity and therapeutic vulnerabilities of human NSCLC.

## Materials and Methods

### Murine cell lines

Murine cell lines from lung adenocarcinomas of conditional cre-lox-cre *Kras*^*G12D*^*Tp53*^*−/−*^*Luc* (KP), *Kras*^*G12D*^*Tp53*^*+/−*^*Lkb1*^*−/−*^*Luc* (KPL) FVB mice that express firefly luciferase were established in Professor David Shackleford’s laboratory. Whole exome sequencing (WES) analysis revealed that KPL cells lost the other allele of *Tp53* upon *in vitro* culture and, therefore, bear a *Kras*^*G12D*^*Tp53*^*−/−*^*Lkb1*^*−/−*^*Luc* genotype. The *Kras*^*G12D*^ LKR-13 line (K) was generously provided by Professor Jonathan Kurie. Each cell line was maintained in culture media (RPMI-1640 medium supplemented with 10% FBS and 1% penicillin/streptomycin) at 37°C in a humidified atmosphere of 5% CO_2_.

### *In vivo* studies

FVB and 129-E mice were purchased from Charles River Laboratories. Tumor cells were implanted in 7-9-week-old mice subcutaneously at optimal doses as indicated in figure legends. Tumor length and width were measured by caliper and the volume calculated by the equation: 0.4*length*width^2. For bioluminescence studies, images were obtained with the IVIS Spectrum imager after intraperitoneal (IP) injection of D-luciferin (150mg/kg). For immunotherapy studies, mice bearing ~50mm^3^ tumors were randomized and treated with 200 μg of anti-PD-1 antibody (BioXcell, Clone RMP1-14) or isotype control via IP injections three times weekly for 4 doses. Mice were housed in pathogen-free facilities at UCLA and all procedures were approved by the UCLA Animal Research Committee.

### Chemical treatment

Cells were seeded in T25 flasks and when ~70% confluency was achieved, culture media was removed and the cells exposed to 100 μg/ml of MNU (Chem Service, NG-17031) in PBS for 45 minutes. After the removal of MNU, cells were washed with PBS twice and fresh culture media was added. Cells were passaged a minimum of three times prior to the subsequent MNU exposure for up to 7 cycles.

### *In vitro* proliferation assay

Cells were plated in culture media in 96-well plates at 1000 cells per well in 8 replicates. Proliferation was measured using ATPlite 1step Luminescence Assay Kit (Perkin Elmer) every 24 hours up to 120 hours. Reading at each time point was normalized to the reading at baseline to control for plating differences.

### *In vitro* IFN-γ stimulation

Cells were seeded in 6-well plates and treated with IFN-γ at 100 ng/ml when 50% confluency was achieved. Cells were harvested 24 hours after stimulation and PD-L1 expression was analyzed by flow cytometry (FACS).

### Tissue preparation

Spleens were mashed with the blunt end of 3 mL syringe on Petri dishes containing 5 ml of PBS, filtered through 70 μM filter, and centrifuged at 1500 rpm for 5 min at 4°C. Cell pellets were resuspended in 5 ml of red blood cell lysis solution (BioLegend) on ice for 5 min followed by the addition of 20 mL of complete media. Cells were filtered through 70 μM filter, centrifuged, washed with PBS, and counted. Murine tumors were harvested, minced with scalpel blades, and digested in 2.5 ml of culture media containing 1 mg/mL of Collagenase IV (Roche) and 50 unit/mL DNase (Sigma) in 15 mL tubes at 37°C with shaking every 10 min. After 45 min, 10 ml of fresh culture media was added and the samples were filtered through 70 μM filter and centrifuged. Red blood cells were lysed as described above, and the cells were washed with PBS, and counted.

### FACS

Single-cell suspension from tumor, spleen, or cell culture were incubated with antibodies for 20 min at 4°C followed by washing with staining buffer (PBS + 2% FBS). Intracellular staining was performed using an eBioscience intracellular kit according to the manufacturer’s protocol. FACS was performed on Attune NxT cytometer (ThermoFisher), and data analyzed by FlowJo software (TreeStar). Details of the flow antibodies utilized are listed in Table S1.

### Genomic profiling

#### Genomic DNA isolation, library preparation, and sequencing

Genomic DNA was extracted from tumor cells (Qiagen, DNeasy blood and tissue kit) for WES. Tail DNA from three FVB and three 129-E mice was included as a normal reference for variant calls. Libraries for WES were prepared using the Kapa Hyper Prep Kit (Roche, KK8504) followed by exome enrichment with SeqCap EZ Share Developer Probe (Roche, 08333025001). Sequencing was performed on Hiseq3000 instrument as 150 bp pair-end runs with the aim of 100x depth at UCLA TCGB Core facility.

#### Sequencing Alignment

Sequence reads were aligned to the mouse genome (mm10) with Burrows-Wheeler Aligner (v 0.7.17), then marked for duplicates and re-calibrated as suggested by Genome Analysis Toolkit (GATK).

#### Variant Calling and Annotation

Strelka2 was utilized to call variants between cell line and the associated normal tail genome^18^. When more than one reference normal genome was available (e.g. FVB mice), variant calls were performed to the individual normal genome independently and kept for down-stream analyses if they were called based on both reference genomes. Finally, a variant was called a mutation if (1) it did not belong to the germline mutation panel, determined from the FVB and 129E tail genomes, (2) it was not supported by any read in the associated normal genome, (3) it was detected by at least 5 reads in cell lines, and (4) its variant allelic frequency (VAF) was > 0.1. The mutations that passed these criteria were then annotated by Ensembl Variant Effect Predictor as nonsynonymous mutations^19^. Mutation with VAF ≥ 0.4 were defined as truncal. Shared mutations with VAF < 0.4 were named as branch. Mutations with VAF < 0.4 which were specific to one cell line and not shared with other members of the family were defined as private.

### Neoantigen prediction

Mutations altering amino acid sequences of the encoded proteins were subjected to MHC-I binding prediction utilizing NetH2pan^20^. Peptides subjected to the algorithm were composed of 8-11 amino acid residues with the mutated amino acid at different locations. A candidate neoantigen had its affinity score ranked < 2^nd^ percentile.

### Statistical analysis

Statistical analyses were performed by GraphPad Prism version 7.04 for Windows. In brief, statistical significance was based on two-tailed non-paired Student’s *t* test for pairwise comparison and two-way ANOVA with Tukey post-test for time-associated comparison among multiple groups. Numerical data was presented as mean ± SEM.

## Results and Discussions

### Establishing mouse models of NSCLC with known driver mutations and varying TMB

Given the low mutational burden in *Kras*-mutant GEMMs of NSCLC, we reasoned that these tumors would be resistant to anti-PD-1 therapy. We screened cell lines established from *Kras*-mutant GEMMs of NSCLC with common co-occuring genomic alterations, namely K, KP, and KPL, with anti-PD-1 antibody in immunocompetent mice and observed primary resistance to immunotherapy across all cell lines except for K, which showed a statistically significant but modest response to PD-1 blockade (Fig 1A, and Fig S1A). Because loss of a functional PD-L1 axis has been implicated in primary resistance to ICIs, we assessed the capacity of IFN-γ to upregulate PD-L1 in K, KP, and KPL cells *in vitro*^21^. IFN-γ stimulation resulted in the upregulation of PD-L1 in all cell lines examined, confirming an intact PD-L1 axis (Fig S1B). To establish cell lines with high TMB, we exposed K, KP and KPL cells to the DNA alkylating agent MNU for various durations (3, 5, and 7 exposures to MNU for 45 minutes each time, designated as 3M, 5M, and 7M, respectively) (Fig 1B). WES of the cell lines exposed to MNU revealed a dose-dependent increase in the number of nonsynonymous mutations across all genomic backgrounds, resulting in higher TMBs that are comparable to human NSCLCs (Fig 1B, and Fig S1C)^12^. In parallel to increased TMB among each isogeneic cell line, we observed increased proportion of mutations with low VAF, suggesting increased tumor heterogeneity (Fig 1C). However, we also detected truncal and branch mutations with high VAF that were shared among Parental, 3M, 5M and 7M cells within each distinct genetic background (Fig 1D).

**Figure 1.**
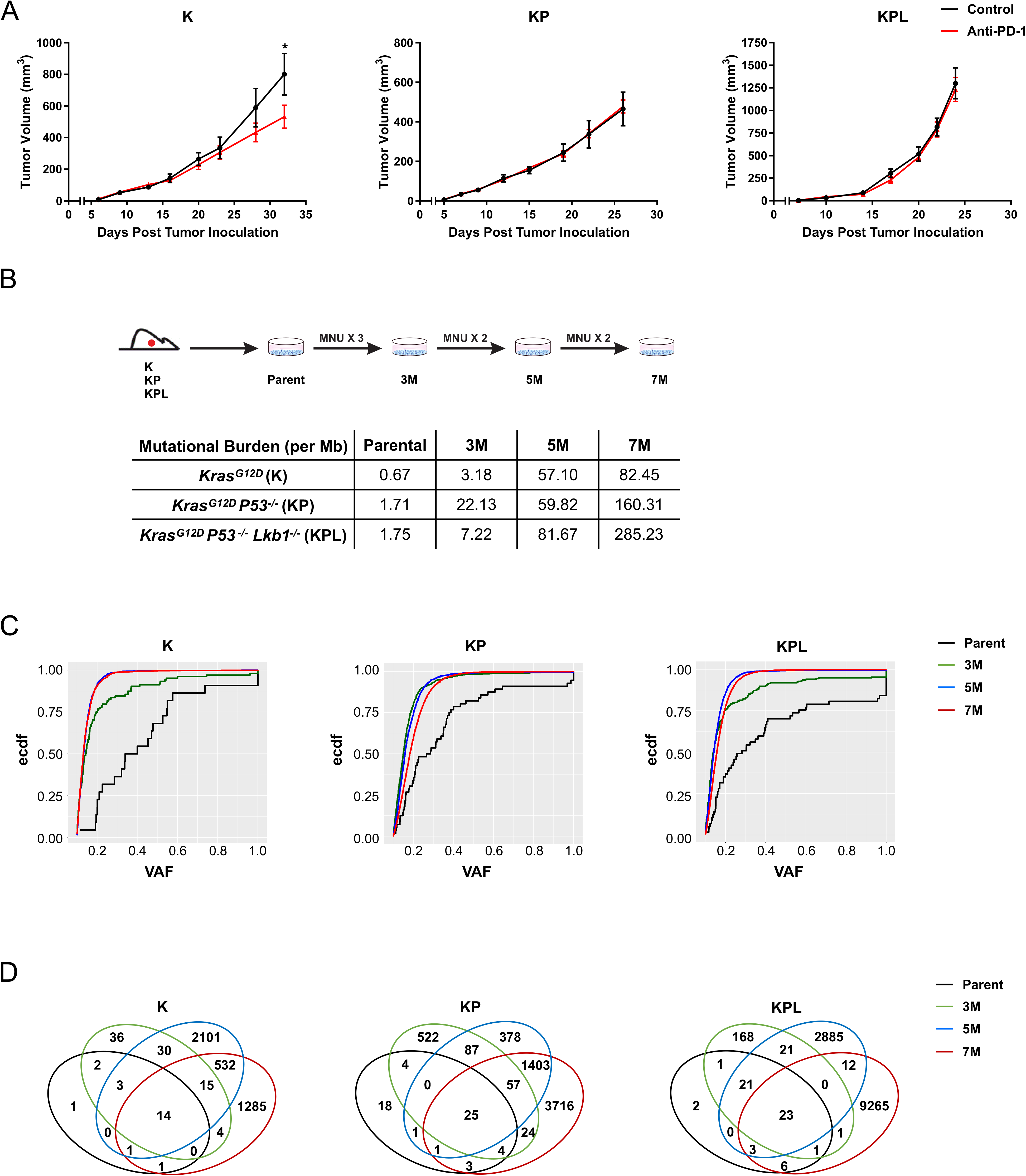
Murine Models of NSCLC with varying mutational burden. **A**) After subcutaneous (SC) tumor inoculation [K (2×10^6^) cells in 129-E mice; KP (8×10^5^) cells in FVB mice; KPL (1×10^5^) cells in FVB mice], mice bearing <50mm^3^ tumors (~days 7) were treated with i) isotype control, Anti-PD-1 (200 μg/dose every 3 days for 4 doses), and tumor growth was measured with caliper. Results are representatives of at least two biological replicates of 6-10 mice per group. **B**) K, KP, and KPL were exposed to 100 μg/ml of MNU for 45 minutes. Cells were passaged prior to additional exposures to MNU for a total of 3, 5, and 7 exposures (3M, 5M, 7M). TMBs obtained from WES are shown in the table. **C)** Empirical cumulative distribution function (ECDF) of the mutations is plotted against VAF as an illustration of tumor heterogeneity within each family of cells. **D**) Venn diagram of shared and private mutations of the K, KP, and KPL isogenic cell lines. *P* values were determined by two-way ANOVA with Tukey post-test. *, *P*<0.05.

### Higher TMB results in decreased tumor growth due to immune rejection

To determine the effect of increased TMB on tumor growth, we screened each family of cell lines with varying TMB in syngeneic models. Across all genomic backgrounds, we observed diminished *in vivo* tumor growth with incremental increases in TMB (Fig 2A). Increasing the number of injected cells led to robust tumor growth of K-3M, KP-3M, KPL-3M and KPL-5M cells, while other lines, namely K-5M, K-7M, KP-5M, KP-7M, KPL-7M, were rejected or displayed diminished growth rates (data not shown). Evaluation of *in vitro* growth rates of cell lines with varying TMB within each family revealed minimal differences (Fig S2A). Therefore, we hypothesized that diminished *in vivo* growth associated with increased TMB was immune-mediated. To test our hypothesis, we evaluated the *in vivo* growth of the K, KP and KPL parental cells, and their associated 7M counterparts in immunocompromised SCID mice which lack T and B cells (Fig 2B). We observed similar tumor growth rates of K-7M and KP-7M compared to their parental counterparts, while KPL-7M showed minor but statistically significant reduction in tumor growth compared to parental KPL cells. These results suggest that the decreased tumor growth rates associated with high TMB in immunocompetent mice are predominantly due to host adaptive immune responses.

**Figure 2.**
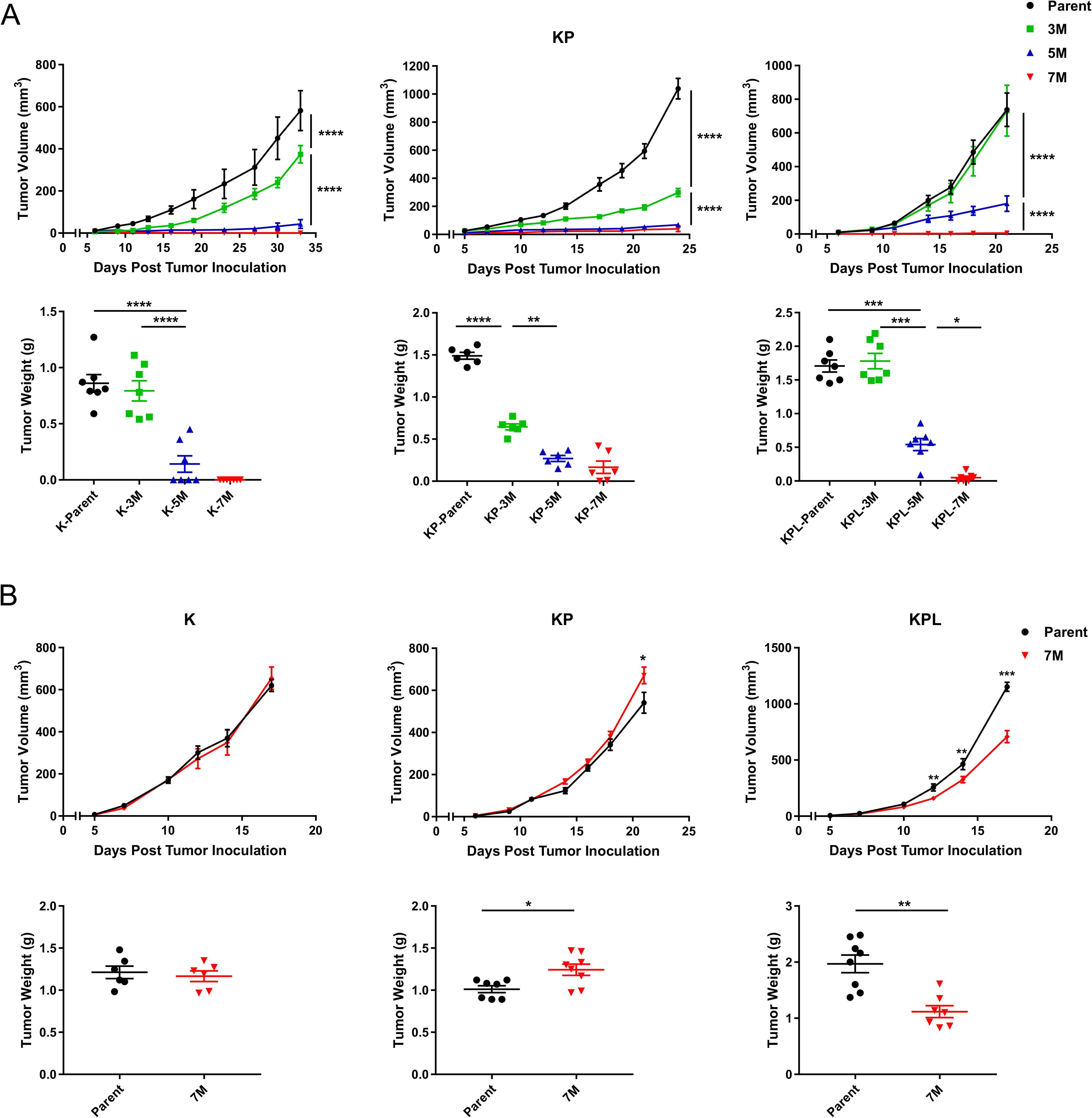
*In vivo* tumor growth in immunocompetent and SCID mice. **A)** Within each family of cells, the Parent, 3M, 5M, and 7M cells were inoculated SC in immunocompetent mice [K (2×10^6^) cells in 129-E mice; KP (8×10^5^) cells in FVB mice; KPL (1×10^5^) cells in FVB mice] and tumor growth was measured with caliper. Growth curves and corresponding tumor weights after euthanasia are presented. **B**) Same as in **A** except Parent and 7M cells were inoculated SC in SCID mice [K (2×10^6^) cells; KP (8×10^5^) cells; and KPL (1×10^5^) cells]. Data are representatives of at least two biological replicates of 6-10 mice per group. *P* values were determined by two-tailed non-paired Student’s *t* test for pairwise comparison and two-way ANOVA with Tukey post-test for time-associated comparison among multiple groups. *, *P*<0.05; **, *P*< 0.01; ***, *P*<0.001; ****, *P*< 0.0001.

### High TMB is associated with increased T cell activation and tumor infiltration

To define the immune responses induced by high TMB, we sought to characterize the immune components of the TME and spleen within each genetic background. Utilizing cells with robust *in vivo* growth, namely K-Parent, K-3M, KP-Parent, KP-3M, KPL-Parent, KPL-3M, and KPL-5M, we first evaluated the lymphoid compartment of the TME. We observed a significant increase in the number of tumor-infiltrating lymphocytes (TILs) and an increase in CD8^+^ to regulatory T (Treg) cell ratio with increased TMB in each genetic background (Fig 3A). Tumor infiltrating CD8^+^ T cells in the tumors with high TMB expressed higher levels of the proliferation marker Ki-67 compared to their respective parental tumors.

**Figure 3.**
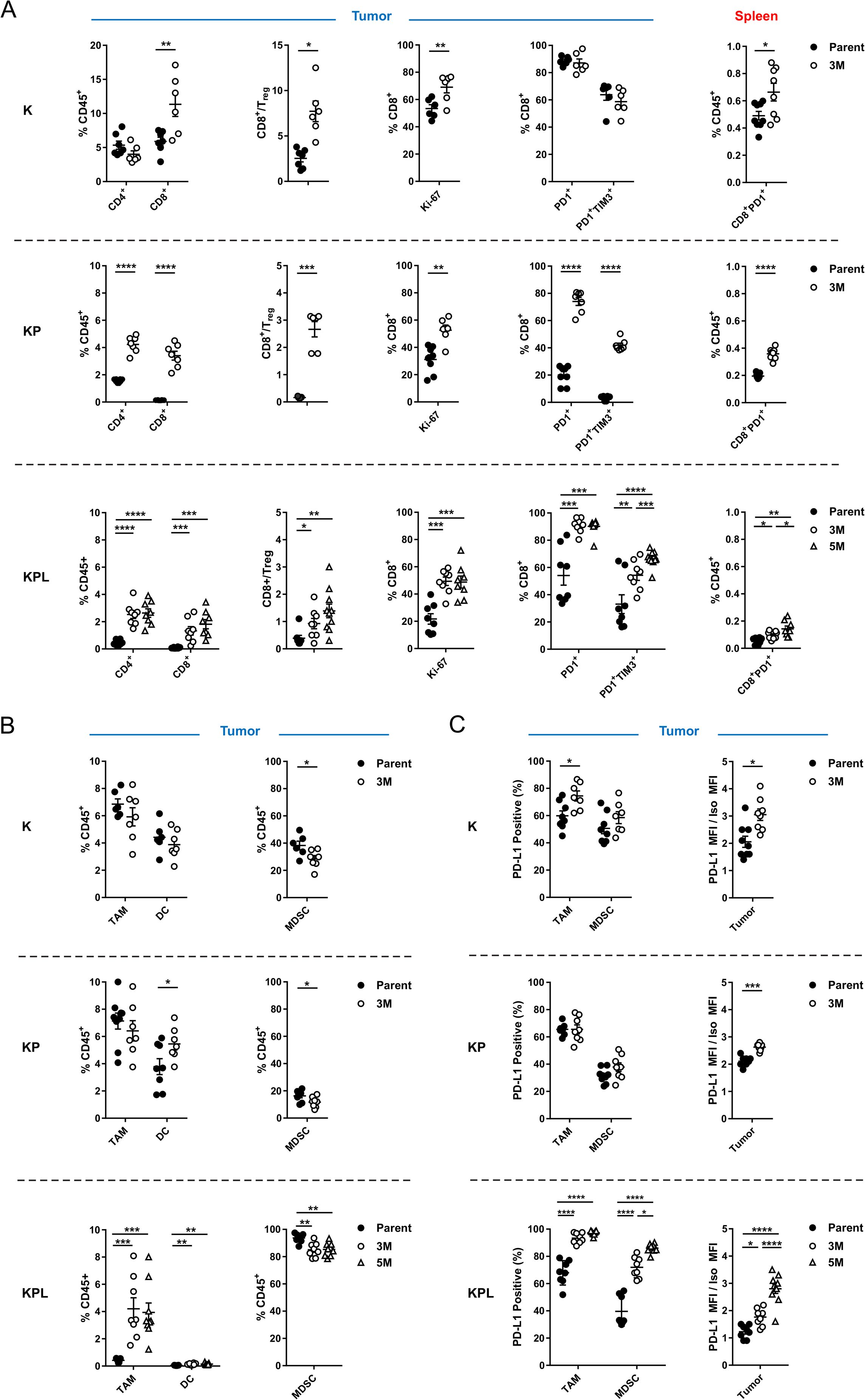
Distinct immune phenotypes of murine models revealed by FACS. On day 14-16 post-tumor inoculation [2×10^6^ K-Parent and K-3M cells in 129-E mice; 8×10^5^ KP-Parent and 2×10^6^ KP-3M cells in FVB mice; 1×10^5^ KPL-Parent, 1.5×10^5^ KPL-3M, and 3×10^5^ KPL-5M cells in FVB mice], tumors and spleens were harvested and analyzed by FACS. **A)** Lymphoid compartment. **B)** Myeloid compartment. **C)** PD-L1 expression. Data are representatives of at least two biological replicates of 6-10 mice per group. *P* values were determined by *P* values were determined by two-tailed non-paired Student’s *t* test. *, *P*<0.05; **, *P*< 0.01; ***, *P*<0.001; ****, *P*< 0.0001.

Next, we evaluated the expression of the early activation/exhaustion marker PD-1 on TILs and observed higher expression of PD-1 on CD8^+^ T cells in KP and KPL tumors with high TMB compared to their parental counterparts (Fig 3A). Studies have revealed that tumor-specific CD8^+^ TILs in human cancers express high levels of PD-1 and that PD-1 expression on CD8^+^ TILs can identify the diverse repertoire of clonally expanded tumor-reactive T cells, therefore our results suggest that increased TMB in the KP and KPL models results in increased tumor-specific CD8^+^ T cell responses^22,23^. In parallel, we also detected increased co-expression of the checkpoint TIM-3, a marker of increased T cell exhaustion with prolonged antigen exposure, on PD-1^+^CD8^+^ T cells in KP and KPL tumors with high TMB compared to their parental counterparts. However, we observed no difference in tumor CD8^+^ T cell exhaustion between K-parent and K-3M tumors. This observation is likely due to increased baseline immunogenicity of the K-parent tumors, which contain high TILs and increased CD8^+^ T cell exhaustion at baseline, and exhibit slow *in vivo* growth with modest sensitivity to anti-PD-1 therapy. In summary, these results support the hypothesis that a higher TMB results in increased tumor-reactive PD-1^+^ T cells within the TME, which become exhausted with persistent antigen stimulation.

Peripheral tumor neoantigen-specific T cells which overlap with clonal tumor-specific TILs have also been identified in circulating PD-1^+^CD8^+^ T cells in melanoma patients^24^ Therefore, we evaluated the expression of PD-1 on CD8^+^ T cells from the spleen of tumor-bearing mice. We observed an increase in splenic PD-1^+^CD8^+^ T cells in mice bearing K, KP, and KPL tumors with high TMB compared to the parental counterparts, including a statistically significant increase in mice bearing KPL-5M tumors compared to those bearing KPL-3M tumors (Fig 3A). This data indicates that increased TMB results in enhanced systemic tumor-specific T cell responses in our murine models.

Next, we evaluated the myeloid compartment of the TME and observed TMB-mediated changes in the immune phenotypes shared across all genetic backgrounds, as well as striking differences specific to KPL cells (Fig 3B, and Fig S3A). Notably, high TMB was associated with a significant increase in the professional antigen-presenting dendritic cells (DCs) in the KP and KPL tumors, with no differences observed in K tumors. We observed no differences in the number of tumor-associated macrophages (TAMs) in K-3M and KP-3M compared to the respective parental tumors, but observed an increase in TAMs in KPL-3M and KPL-5M tumors compared to KPL-Parent. Next, we evaluated changes in the MDSCs and found that high TMB was associated with decreased MDSCs across all genetic backgrounds. Notably, KPL tumors contained a significantly higher percentage of MDSCs (over 80% of CD45^+^ cells), which predominantly expressed the neutrophil marker Ly6G (Fig 3B, and S3B). This phenotype is consistent with prior studies in *KRAS*-mutant murine and human NSCLC where LKB1 loss has been associated with a T cell-suppressed and neutrophil-enriched TME^25,26^.

We further evaluated changes in the PD-L1 expression by tumors and the myeloid cells in the TME (Fig 3C). We observed an increase in PD-L1 expression on TAMs in K and KPL tumors with increased TMB compared to their parental counterparts but no difference between KP-3M and KP-Parent. Furthermore, we observed increased PD-L1 expression on MDSCs in the KPL tumors with higher TMB, with the greatest expression observed in KPL-5M, but no difference was detected within the K and KP genetic background. In addition, we observed increased PD-L1 expression on tumors with increased TMB. Taken together, the observed overall trend of increased PD-L1 expression associated with increasing TMB implies amplified adaptive immune resistance within the TME in response to increased anti-tumor responses.

### High TMB is associated with enhanced efficacy of anti-PD-1

Given that high TMB in our murine models was associated with enhanced local and systemic T cell activation and increased PD-L1 expression within the TME, we hypothesized that high TMB could sensitize tumors to PD-1 blockade. To test this hypothesis, we evaluated the efficacy of PD-1 blockade in K-3M, KP-3M and KPL-3M cells (Fig 4A and Fig S4A). Anti-PD-1 therapy resulted in robust anti-tumor responses with an eradication of 33% of K-3M tumors. Similarly, 44% of KP-3M tumors were rejected and others stabilized in response to anti-PD-1. In contrast, anti-PD-1 efficacy was modest in KPL-3M tumors where PD-1 blockade resulted in reduced tumor growth without a complete rejection. This result is in agreement with the recent findings in human *KRAS*-mutant lung adenocarcinoma where LKB1 loss was shown to be a major driver of primary resistance to PD-1 blockade^26^. We next assessed the efficacy of PD-1 blockade in mice bearing KPL-5M and observed significant anti-tumor responses with the rejection of approximately 50% of tumors (Fig 4B). This data suggests that the increased TMB of KPL-5M tumors could overcome the immunosuppressed TME and enhance responses to PD-1 blockade. This is in agreement with our immunophenotyping results of the KPL family of tumors where mice bearing KPL-5M tumors possessed the highest number of local and systemic activated PD1^+^ CD8^+^ T cells (Fig 3), which have been shown to contain pools of tumor noantigen-specific T cells that can be reinvigorated following PD-1 blockade^10,27^. Next, we challenged the mice that were cured of KPL-5M cancer upon anti-PD-1 therapy with KPL-5M cells and observed an initial tumor growth followed by spontaneous rejection of all tumors (Fig S4B). These results indicate the establishment of systemic anti-tumor immunity in response to PD-1 blockade in mice bearing KPL-5M tumors.

**Figure 4.**
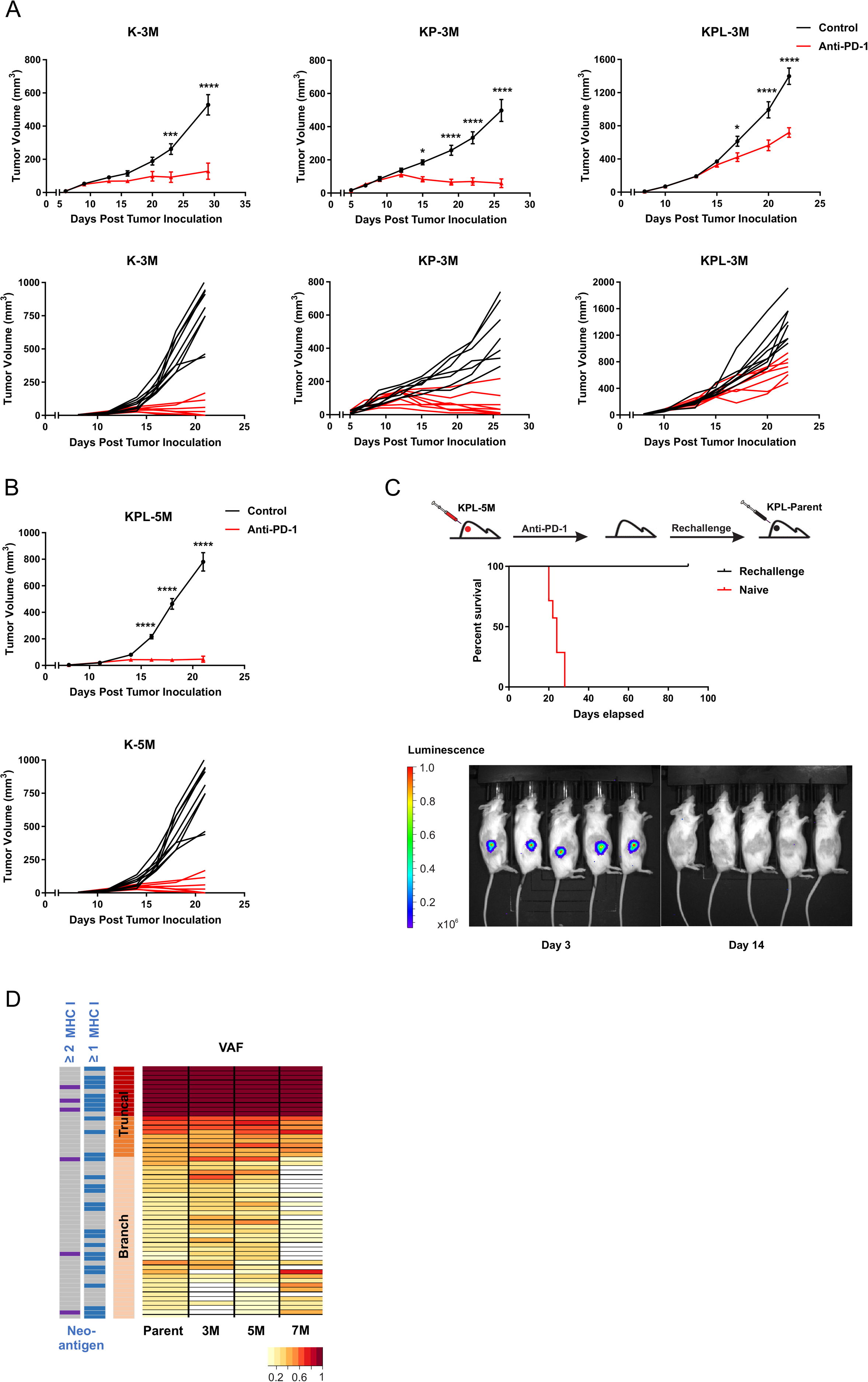
High TMB results in increased efficacy of anti-PD-1 therapy. **A)** After SC tumor inoculation [K-3M (2×10^6^) cells in 129-E mice; KP-3M (2×10^6^) cells in FVB mice; KPL-3M (1.5×10^5^) cells in FVB mice], mice bearing <50mm^3^ tumors (~days 7) were treated with i) isotype control, ii) Anti-PD-1 (200 μg/dose every 3 days for 4 doses), and tumor growth was measured with caliper. Results are representatives of at least two biological replicates of 6-10 mice per group. **B)** Same experimental design as **A** except that KPL-5M (3×10^5^) cells were utilized for SC tumor inoculation. **C)** FVB naïve mice and FVB mice that previously eradicated KPL-5M tumors in response to PD-1 blockade were inoculated SC with KPL-Parent (2×10^5^) cells and tumor growth was measured with bioluminescence imaging on day 3 and day 14. Survival curve is presented. Data is representatives of two biological replicates of 5-6 mice per group. **D)** Representation of the frequency of the mutations in the KPL-Parent tumors shared by KPL-3M, KPL-5M, or KPL-7M. Predicted neoantigens based on MHC-I binding avidity are also presented. *P* values were determined by two-way ANOVA with Tukey post-test. *, *P*<0.05; **, *P*< 0.01; ***, *P*<0.001; ****, *P*< 0.0001.

Given that KPL-5M shares 47 truncal and branch mutations with KPL-Parent (Fig 1C), we assessed whether the mice that had eradicated the KPL-5M tumors after anti-PD-1 treatment could reject the parental tumors, by inoculating the mice with KPL-Parent 3 months after the initial rejection (Fig 4C). Indeed, all of the mice eliminated the KPL-Parent tumors after an initial growth, while the naïve control mice succumbed to implantation of KPL-Parent tumors in less than 30 days. Computational analysis of putative neoantigens in the KPL-Parent revealed 11 truncal and 17 branch neoantigens which were shared with KPL-3M, KPL-5M or KPL-7M (Fig 4D, and Fig S4C). These results indicate the presence of tumor-specific memory T cells against shared neoantigen(s) between KPL-5M and KPL-Parent tumors in anti-PD-1 treated mice that had eradicated KPL-5M tumors. Yet, our data indicates that T cell responses against these shared neoantigen(s) are absent or not sufficient to mount KPL-Parent tumor rejection in naïve mice treated with PD-1 blockade (Fig 1A), which is possibly a result of profound immunosuppression in the TME of KPL-Parent tumors (Fig 3). In contrast, eradication of KPL-P tumors in the rechallenge experiments is likely predominantly mediated by memory T cells, which can generate a rapid recall response to secondary tumor-neoantigen(s) challenge. The presence of shared neoantigen(s) in these isogenic cell lines with varying TMB provides a unique opportunity to investigate immune responses against truncal and branch mutations in the context of TMB-associated changes in the TME.

In summary, herein we report novel *Kras*-mutant murine models of NSCLC bearing common genetic alterations of the disease and physiologically relevant TMB that recapitulate the genetic complexity of *KRAS*-mutant NSCLC in human. Similar to human NSCLC, these preclinical models possess intratumoral heterogeneity as well as truncal and branch neoantigens that can elicit adaptive immune responses. Moreover, these *Kras*-mutant murine models with co-occurring oncogenic mutations have distinct TMEs, and recapitulate the therapeutic vulnerabilities to PD-1 blockade observed in human *KRAS*-mutant NSCLC. We anticipate that these novel immunogenic murine models of NSCLC will serve as relevant preclinical platforms for mechanistic interrogation of neoantigen evolution and tumor-specific immune responses to cancer immunotherapy and facilitate the development of novel therapies.

