## Supplemental table and figures for "Novel *Kras*-mutant murine models of non-small cell lung cancer possessing co-occurring oncogenic mutations and increased tumor mutational burden"

Table S1. FACS antibodies.

| Immune Marker | Fluorophore | Clone |
|---|---|---|
| CD45 | perCP/Cy5.5 | 30-F11 |
| CD3 | AF700 | 17A2 |
| CD8 | BV421 | 53-6.7 |
| CD4 | BV650 | GK1.5 |
| CD25 | APC | PC61 |
| FOXP3 | PE | 150D |
| PD1 | PE-Cy7 | 29F.1A12 |
| Tim3 | PE | RMT3-23 |
| Ki67 | BV605 | 16A8 |
| CD11c | FITC | HL3 |
| IA/IE | BV650 | M5/114.15.2 |
| CD11b | AF700 | M1/70 |
| Ly6C | APC | HK1.4 |
| Ly6G | PE | 1A8 |
| CD64 | PE-Cy7 | X54-5/7.1 |
| PD-L1 | BV421 | 29E.2A3 |
| Zombie NIR | | |

Fig S1

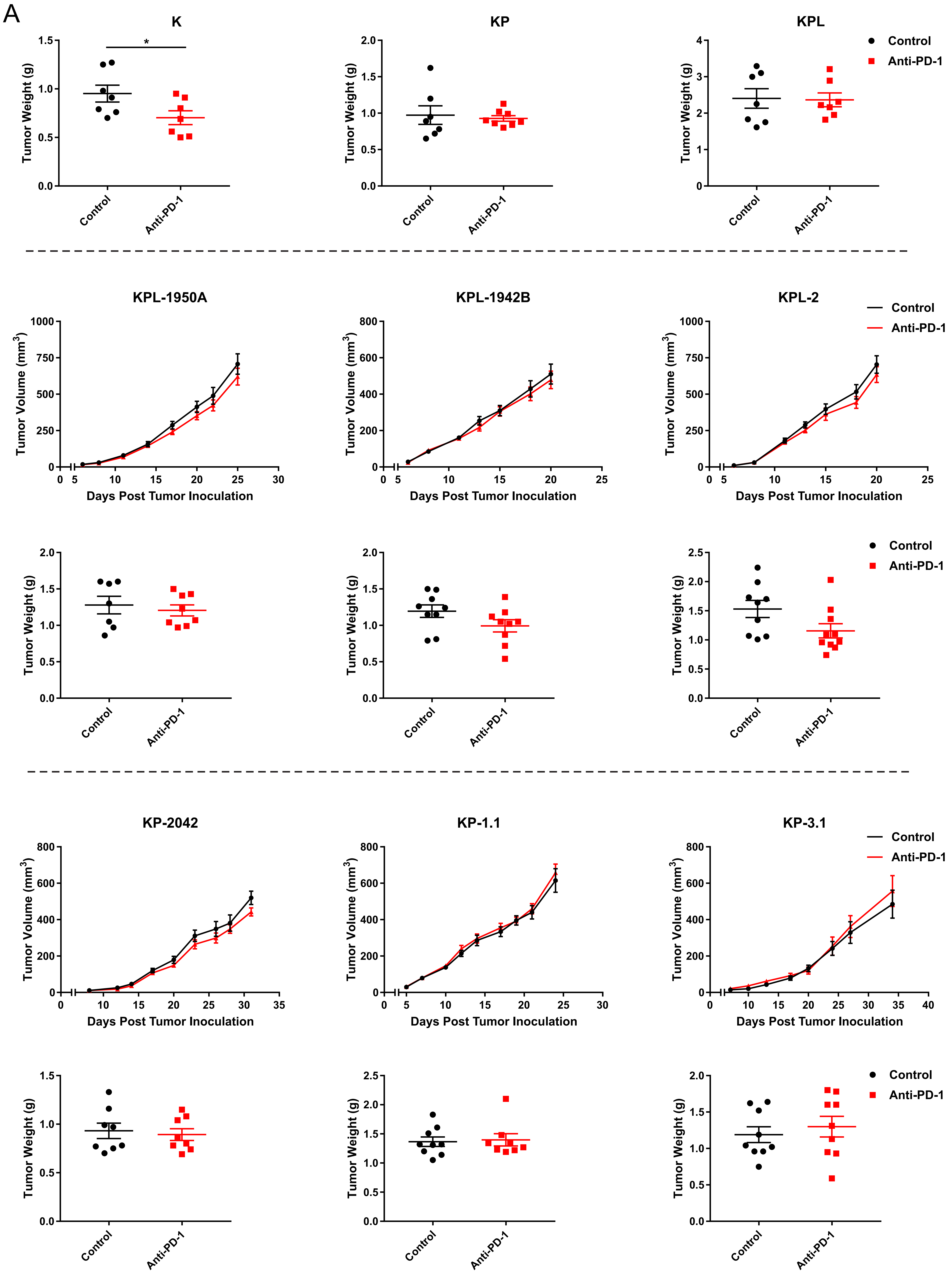
A
K
KP
KPL
Tumor Weight (g)
Control
Anti-PD-1
*
KPL-1950A
KPL-1942B
KPL-2
Tumor Volume (mm3)
Days Post Tumor Inoculation
KP-2042
KP-1.1
KP-3.1

Fig S1

B

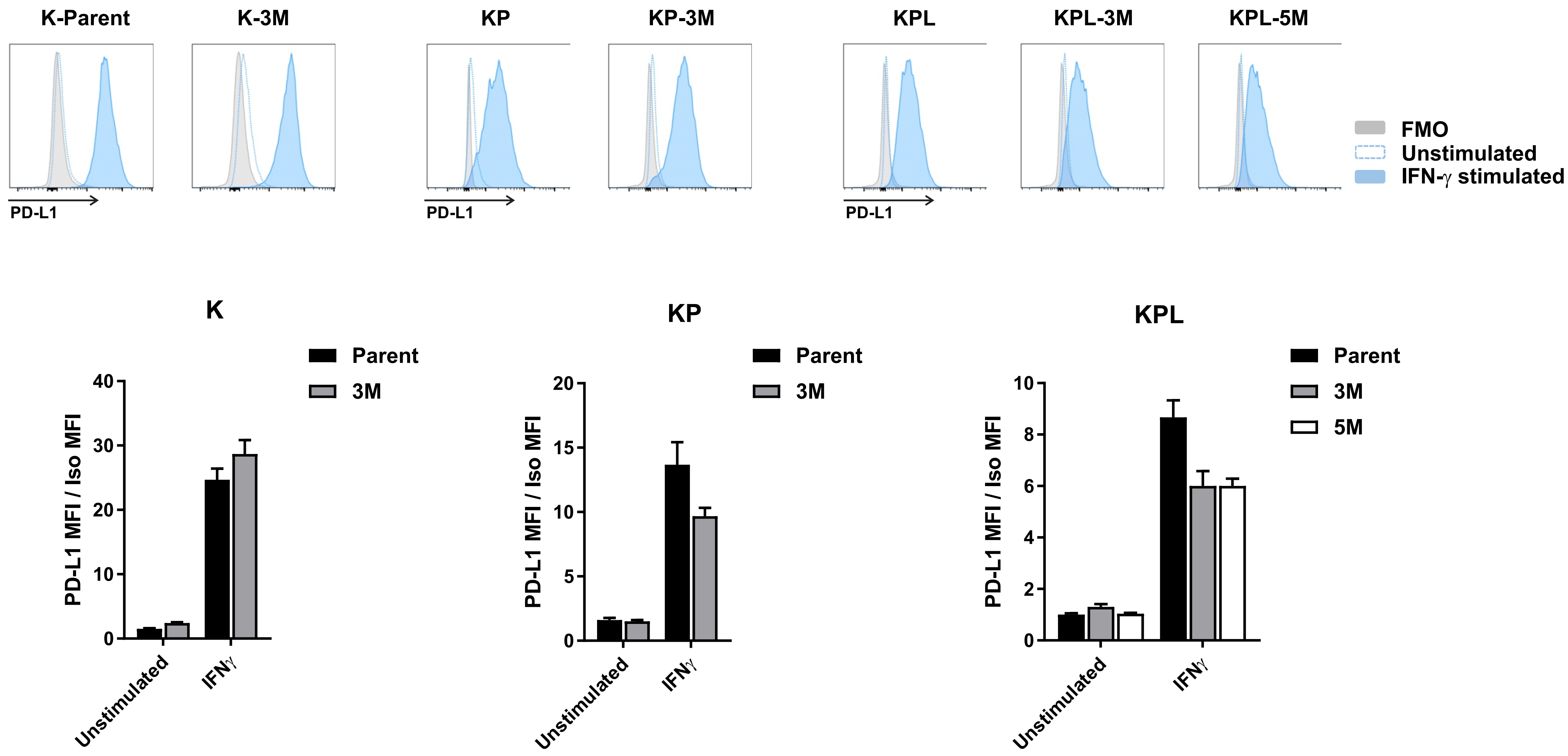


C

| Mutational load (SNV) | Parental | 3M | 5M | 7M |
|---|---|---|---|---|
| ***Kras*$^{G12D}$ (K)** | 22 | 104 | 1852 | 2696 |
| ***Kras*$^{G12D}$ *P53*$^{-/-}$ (KP)** | 56 | 723 | 1952 | 5233 |
| ***Kras*$^{G12D}$ *P53*$^{-/-}$ *Lkb1*$^{-/-}$ (KPL)** | 57 | 236 | 2665 | 9311 |

**Figure S1.**

**A)** K ($2x10^6$) cells were SC inoculated in 129-E mice, and KP and KPL cells were SC inoculated in FVB mice [ KP ($8x10^5$) cells; KPL ($1x10^5$) cells; KPL-1950A ($3x10^5$) cells; KPL-1942B ($1x10^6$) cells; KPL-2 ($1x10^6$) cells; KP-2042 ($1.5x10^5$) cells; KP-1.1 ($2x10^6$) cells; KP-3.1 ($2x10^6$) cells] and tumor growth was measured with caliper. Growth curves and corresponding tumor weights after euthanasia are presented. Data are representatives of at least two biological replicates of 6-10 mice per group. **B)** *In vitro* PD-L1 expression of K, KP and KPL cells (triplicates) at baseline and after stimulation with IFN-$\gamma$ at 100 ng/ml for 24 hours. **C)** Tumor mutational load of K, KP and KPL family of cells. *P* values were determined by two-tailed non-paired Student's *t* test for pairwise comparison and two-way ANOVA with Tukey post-test for time-associated comparison among multiple groups. *, $P<0.05$.

Fig S2

A

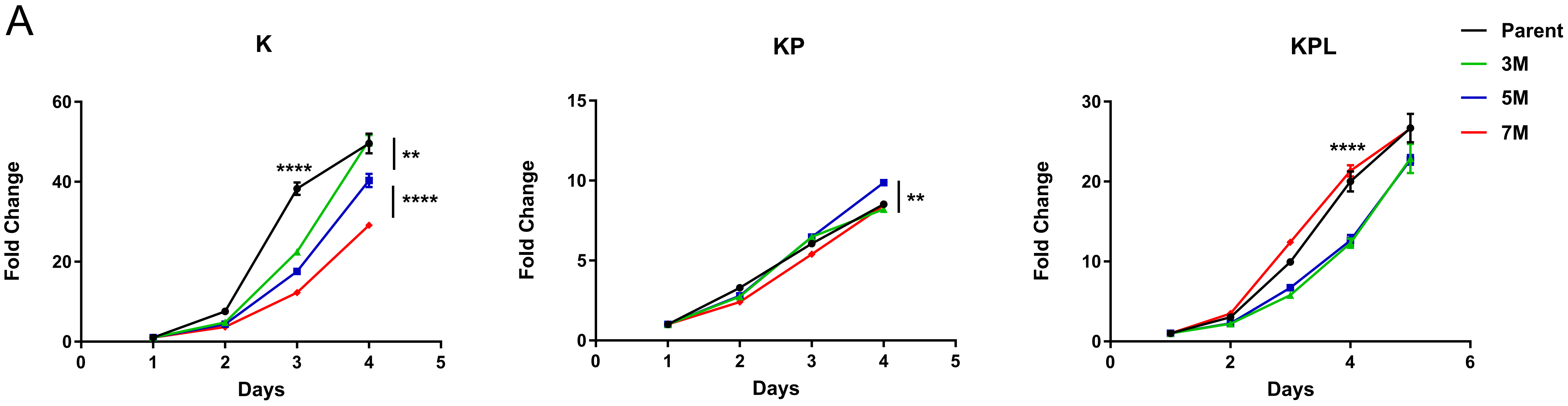

**Figure S2.**

**A)** *In vitro* proliferation of K, KP and KPL family of cells by ATPlite in 8 replicates. Data at each time point is normalized to the reading at baseline to control for plating differences. *P* values were determined by two-way ANOVA with Tukey post-test. *, $P$<0.05; **, $P$< 0.01; ***, $P$<0.001; ****, $P$< 0.0001.

Fig S3

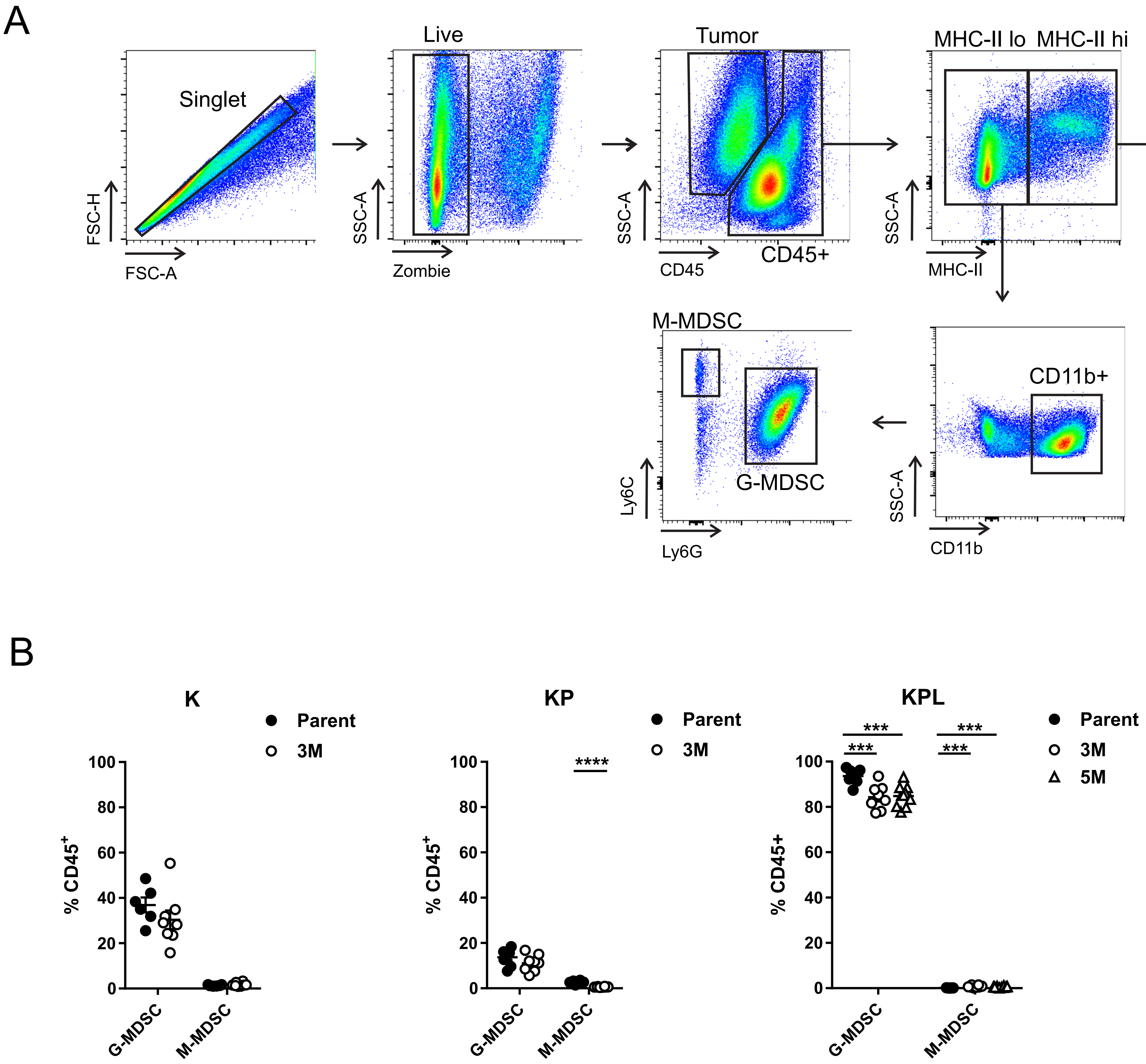
A
Singlet
FSC-H
FSC-A
Live
SSC-A
Zombie
Tumor
CD45
CD45+
MHC-II lo
MHC-II hi
MHC-II
CD64-
TAM
CD11b
CD64
M-MDSC
G-MDSC
Ly6C
Ly6G
CD11b+
CD11b
DC
CD11c
B
K
Parent
3M
% CD45+
G-MDSC
M-MDSC
KP
****
KPL
***
5M

**Figure S3.**

**A)** Flow gating strategy for the myeloid compartment. **B)** On day 14-16 post-tumor inoculation [$2x10^6$ K-Parent and K-3M cells in 129-E mice; $8x10^5$ KP-Parent and $2x10^6$ KP-3M cells in FVB mice; $1x10^5$ KPL-Parent, $1.5x10^5$ KPL-3M, and $3x10^5$ KPL-5M cells in FVB mice], tumors were harvested and analyzed by FACS. Percentage of granulocytic-MDSC (G-MDSC) and monocytic-MDSC (M-MDSC) within the $CD45^+$ compartment is presented. Data are representatives of at least two biological replicates of 6-10 mice per group. *P* values were determined by two-tailed non-paired Student's *t* test. *, *P*<0.05; **, *P*< 0.01; ***, *P*<0.001; ****, *P*< 0.0001.

Fig S4

A

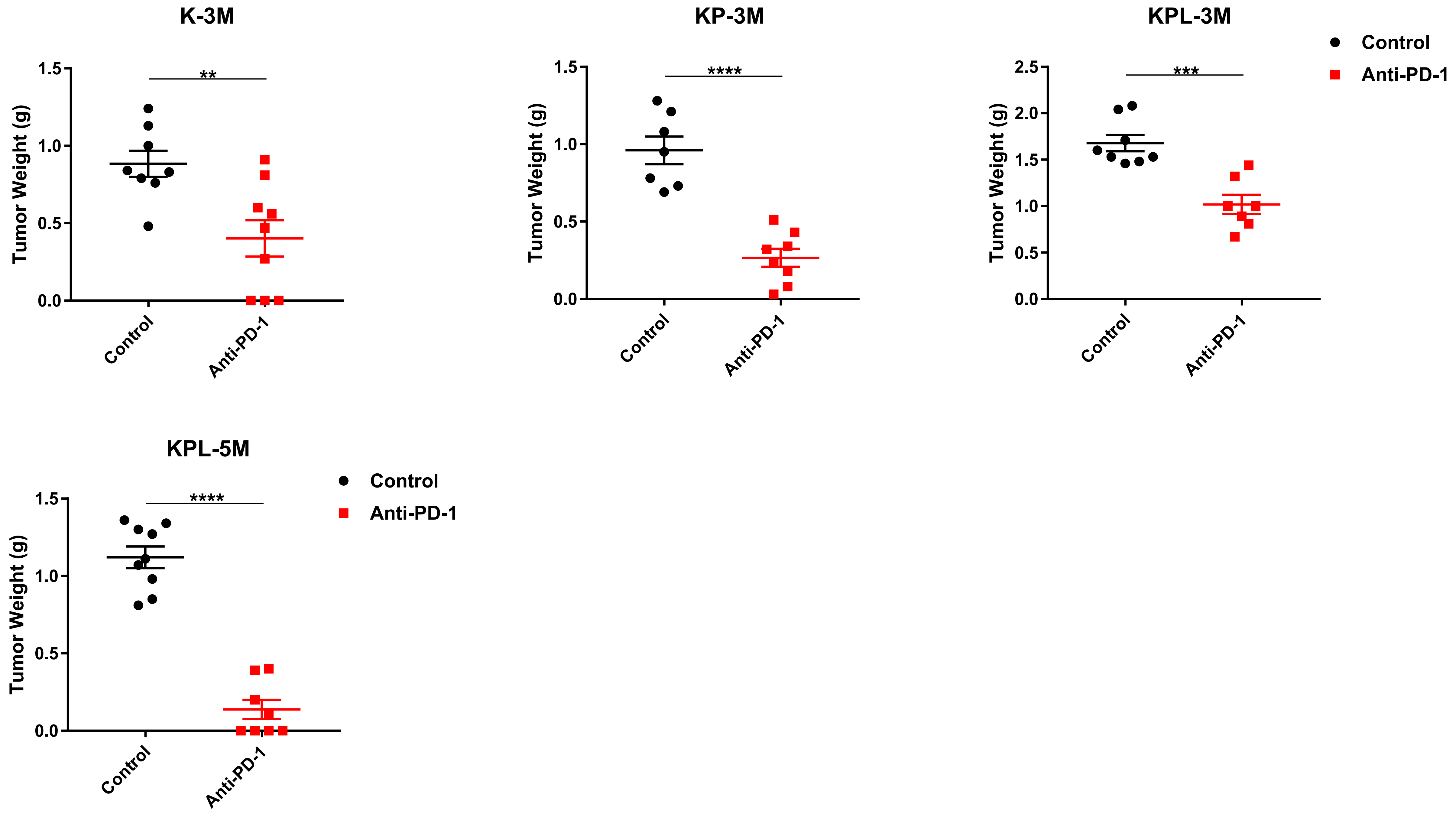
K-3M
KP-3M
KPL-3M
KPL-5M
Tumor Weight (g)
**
****
***
****
Control
Anti-PD-1
Control
Anti-PD-1

B

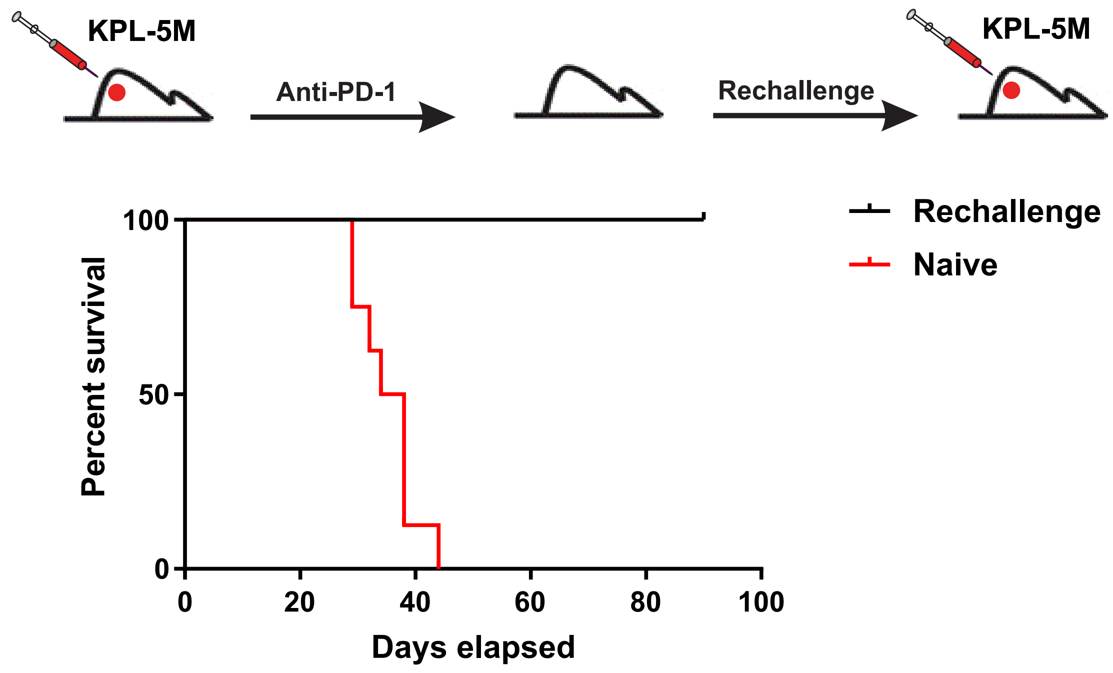
KPL-5M
Anti-PD-1
Rechallenge
KPL-5M
Rechallenge
Naive
Percent survival
Days elapsed
0
20
40
60
80
100

C

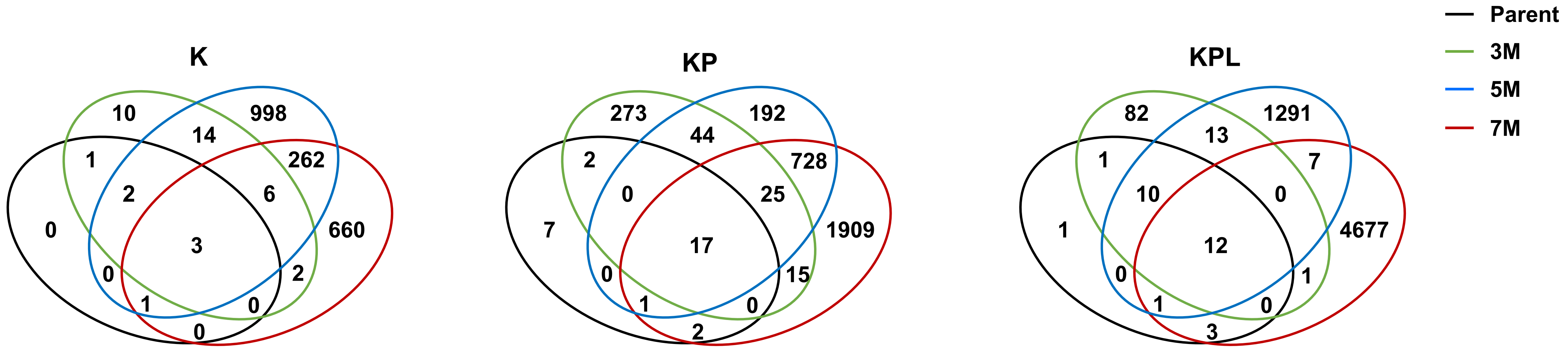
K
KP
KPL
Parent
3M
5M
7M
10
998
14
1
262
2
6
0
3
660
0
2
1
0
0
273
192
44
2
728
0
25
7
17
1909
0
15
1
0
2
82
1291
13
1
7
10
0
1
12
4677
0
1
1
0
3

**Figure S4.**

**A)** After SC tumor inoculation [K-3M ($2x10^6$) cells in 129-E mice; KP-3M ($2x10^6$) cells in FVB mice; KPL-3M ($1.5x10^5$) cells in FVB mice; KPL-5M ($3x10^5$) cells in FVB mice], mice bearing <$50mm^3$ tumors (~days 7) were treated with i) isotype control, ii) Anti-PD-1 (200 µg/dose every 3 days for 4 doses). Tumor weights at the time of necropsy are presented. **B)** FVB naïve mice and mice that previously eradicated KPL-5M tumors in response to PD-1 blockade were inoculated SC with KPL-5M ($3x10^5$). Survival curve is presented. Data is representatives of two biological replicates of 5-6 mice per group. **C**) Venn diagram of predicted MHC-I neoantigens for K, KP, and KPL family of cells. *P* values were determined by two-tailed non-paired Student's *t* test. *, $P$<0.05; **, $P$< 0.01; ***, $P$<0.001; ****, $P$< 0.0001.
